# Fennec - Functional Exploration of Natural Networks and Ecological Communities

**DOI:** 10.1101/194308

**Authors:** Markus J. Ankenbrand, Sonja Hohlfeld, Lorenz Weber, Frank Förster, Alexander Keller

## Abstract

1. Species composition assessment of ecological communities and networks is an important aspect of biodiversity research. Yet often ecological traits of organisms in a community are more informative than scientific names only. Furthermore, other properties like threat status, invasiveness, or human usage are relevant for many studies, but cannot be evaluated from taxonomy alone. Despite public databases collecting such information, it is still a tedious manual task to enrich community analyses with such, especially for large-scaled data.
2. Thus we aimed to develop a public and free tool that eases bulk trait mapping of community data in a web browser, implemented with current standard web and database technologies.
3. Here we present the Fennec, a workbench that eases the process of mapping publicly available trait data to the user’s communities in an automated process. Usage is either by a local self-hosted or a public instance (https://fennec.molecular.eco) covering exemplary traits. Alongside the software we also provide usage and hosting documentation as well as online tutorials.
4. The Fennec aims to motivate public trait data submission and its reuse in meta-analyses. Further, it is an open-source development project with the code freely available to use and open for community contributions (https://github.com/molbiodiv/fennec).

## German Abstract

1. Die Untersuchung der taxonomischen Zusammensetzung ökologischer Gemeinschaften und Netzwerke ist ein zentraler Aspekt in der Biodiversitätsforschung. Eine ausschließlich taxonomische Klassifizierung ist jedoch in vielen Fällen unzureichend, um Zusammenhänge erklären zu können. Hierzu wird vielmehr zusätzliches Wissen über ökologische, morphologische, funktionale und weitere Eigenschaften der Organsimen benötigt. Solche Informationen werden in öffentliche Datenbanken gesammelt, aber es ist bislang ein mühsamer manueller Prozess, gesamte Artgemeinschaften mit diesen Daten anzureichern, insbesondere in Zeiten immer größer werdender Untersuchungen.
2. Daher haben wir ein öffentliches und frei nutzbares Werkzeug entwickelt, welches diesen Kombinationsschritt vereinfacht. Durch den Einsatz von modernen Web- und Datenbank Technologien, konnten wir dieses plattform-unabhängig gestalten.
3. Wir präsentieren hier die webbasierte Anwendung Fennec, welche in einem automatisierten Prozess zu taxonomischen Artengemeinschaften des Nutzers öffentlich oder privat verfügbare Informationen der Organismen hinzufügt. Fennec kann einfach in einer selbstbetriebenen Instanzen ausgeführt werden. Alternativ kann auch eine öffentlichen Instanz (https://fennec.molecular.eco) genutzt werden, die zusätzlich Beispieldaten enthält. Neben einer ausführlichen Dokumentation stellen wir auch Anleitungen als Einstiegshilfe zur Verfügung.
4. Der Fennec soll Wissenschaftler ermutigen Daten zu Organismen in öffentlichen Datenbanken zu hinterlegen, für Meta-Analysen zu nutzen und für andere Wissenschaftler nutzbar zu machen. Das gesamte Projekt folgt den Richtlinien zu quelloffener Entwicklung. Damit ist der Quelltext frei verfügbar und Fennec selbst offen für Beiträge anderer Entwickler (https://github.com/molbiodiv/fennec).

## Introduction

An important task in biodiversity research is the analysis of species composition of ecological communities and networks. This can be done using traditional methods and more recently also with analytical methods designed for large scale sample processing, like DNA metabarcoding (Keller, Danner, et al., 2015) or automated image analysis (Oteros et al., 2015) producing data volumes hard to cope with manually. Therefore, tools for automated taxonomic identification have been developed, e.g. QIIME (Caporaso et al., 2010) and mothur (Schloss et al., 2009).

Ecological or socio-economical hypotheses can however often not be evaluated by looking at taxonomy alone (Junker et al., 2015; Xu et al., 2014), but meta-data for each community-member (e.g. life-history, size or threat-status) becomes important. In microbial ecology, development of tools has already been initiated that aim to automatically map taxonomy information to functional traits, mostly through mapping on known genomes (Aßhauer et al., 2015; Edgar, 2017; Keller, Horn, et al., 2014; Langille et al., 2013). To our knowledge, it remains to date a manual effort to enrich eukaryotic communities similarly with trait meta-data, although such information is already publicly available.

Databases have been developed that provide trait information for eukaryotes and prokaryotes, e.g. LEDA Traitbase (Kleyer et al., 2008), TRY (Kattge et al., 2011), and BacDive or ProTraits for microbial traits (Brbić et al., 2016; Söhngen et al., 2016). On a higher level, TraitBank (Parr, Wilson, Leary, et al., 2014; Parr, Wilson, Schulz, et al., 2014) aggregates this information from different sources. These sources are of course far from complete, yet the existing data is already highly informative.

When using these databases though, currently these trait data have to be manually searched and mapped to the community, which can comprise hundreds or even thousands of taxa. This is impractical, trait data should be accessible with automatic batch annotation procedures, not only single manual requests. This becomes crucial if many taxa are regarded in an ecological community or multiple traits are to be analyzed simultaneously. Furthermore, tools for visualization and interactive analysis of community data like Phinch and phyloseq are currently limited with respect to traits as they only accept taxonomy as metadata for operational taxonomic units (OTUs) (Bik et al., 2014; McMurdie et al., 2013). Here we present the Fennec, a web-based workbench that helps researchers enrich their taxonomy-based community and network interaction tables with relevant traits for their research questions. The workbench includes also basic tools for interactive visualization and analysis of trait data.

A public instance is hosted at https://fennec.molecular.eco, which currently holds roughly 1.7 million organisms and 224 thousand trait entries (referring to 73.5 thousand distinct taxa) gathered from various sources. User-provided community and network data can be readily mapped to these traits.

An alternative way to use the Fennec is to download and host a local instance for user and private traits alongside the ones in the public database. It is freely available at https://github.com/molbiodiv/fennec as an open-source project and we encourage community development and re-use of the code.

Online documentation is available on how to use the public instance and set up own instances (https://fennec.readthedocs.io). This documentation also includes a case study in pollination ecology to demonstrate how the Fennec can be used to investigate specialization of solitary bees on plants with specific growth habit, distribution of invasive taxa as well as occurence of endangered organisms.

## Material and Methods

### General Structure

A Fennec instance consists of a web server and one or multiple databases (Figure 1), which store organism and corresponding trait information. This data can originate from various sources and is imported by the instance administrator. Users can interact with Fennec exploratively or with relation to a specific project using their web browser. The Fennec provides mechanisms to browse and search the contents of its databases including traits (Figure 2A) and organisms. For organisms detailed pages with texts and images from EOL (Parr, Wilson, Leary, et al., 2014) as well as traits and taxonomic information are provided (Figure 2B). Logged in users have access to their individual project section. Users can upload their ecological community/network matrices, match these against available trait data for contained organisms and select traits of interest to the project. The enriched projects can be interactively visualized using the built-in version of Phinch (Bik et al., 2014) or exported for analysis with external tools. (Figure 2C and D).

**Figure 1:**
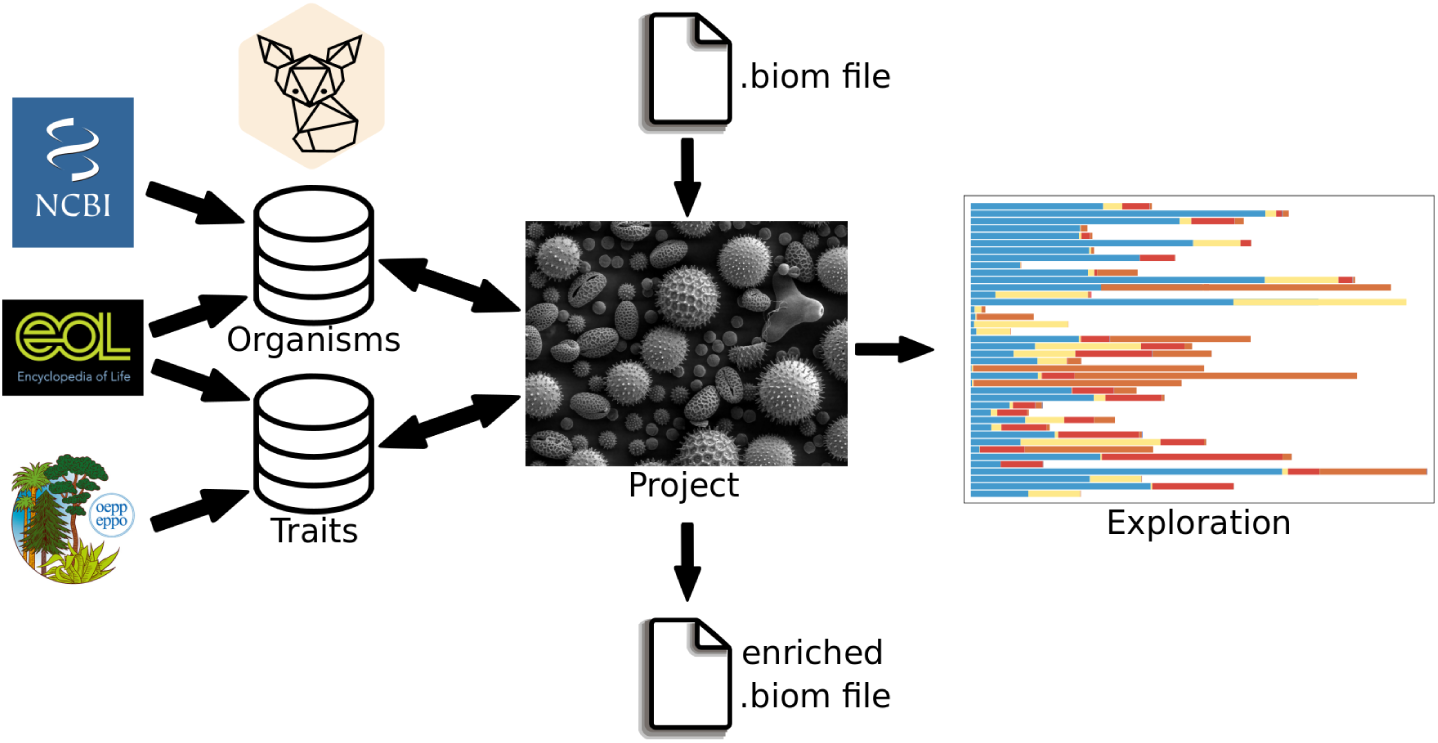
General structure of Fennec. Organisms and traits from different sources are stored by the administrator. The user imports a community project e.g. in biom format. Organisms in the community are mapped against those in the database and enriched with traits. The trait composition can be interactively explored or exported as enriched projects for downstream software.

**Figure 2:**
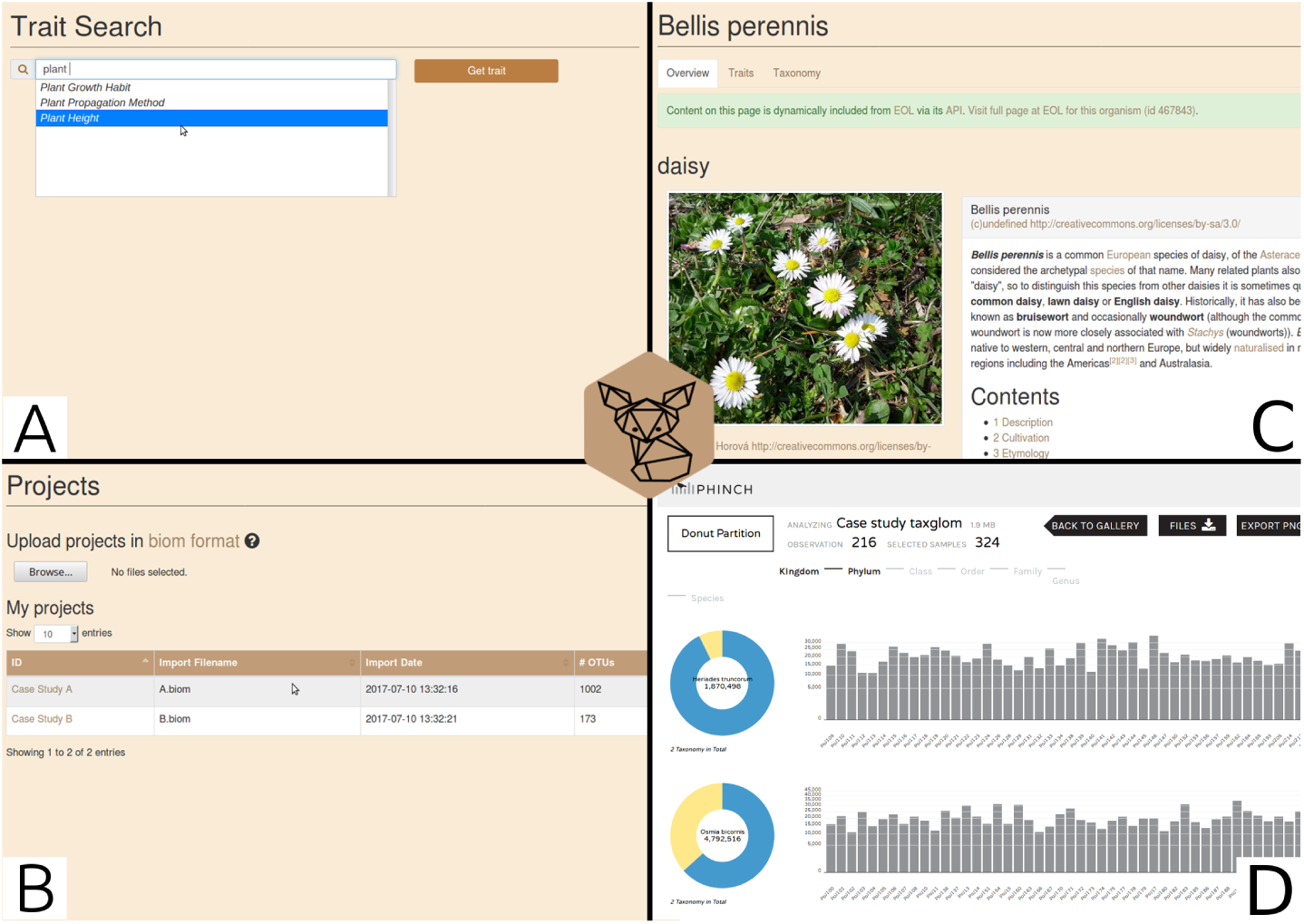
User interface of Fennec. A: Explore traits stored in the database via the search. B: Upload community data, manage and analyze them. C: Get dynamic content for organisms. D: Visualize data using the “Donut Partition Chart”.

### Code Implementation

The Fennec is a web application developed in PHP (https://php.net/) using Symfony (https://symfony.com/) and a JavaScript (ES6) front-end. Server side functionality is bundled in modular web services called via AJAX requests. Layout and interactivity are provided by multiple well-established libraries including bootstrap (https://getbootstrap.com/), jQuery-ui (https://jqueryui.com/), react (https://facebook.github.io/react/), lodash (https://lodash.com/), datatables (https://datatables.net/), and plotly.js (https://plot.ly/javascript/).

Code quality is ensured by tests and strategies for continuous integration. Data is stored in a PostgreSQL (https://www.postgresql.org/) database and accessed via the doctrine object-relational mapper (http://www.doctrine-project.org/). User provided data is uploaded and stored in BIOM format (version 1.0) (McDonald et al., 2012) using the biojs-io-biom library (Ankenbrand et al., 2017).

## Results

### Accessibility

There are three ways to use the Fennec workbench:

1. Public Instance: We have set up a public instance of the Fennec available at https://fennec.molecular.eco. Its database currently hosts trait data related to pollination and microbiomes from various sources. The database is subject to constant further extension with more traits, yet our main goal is to maintain high quality of the data available here. User-provided network and community data is private by default, requiring the user to authenticate using a Fennec or GitHub account.
2. Local Instances: All program code is open-source (MIT License) and freely available at the public repository: https://github.com/molbiodiv/fennec. Alongside the code, ready-to-use (pre-configured) docker containers are available to be run in a virtualization environment (https://www.docker.com). Local instances can be restricted in accessibility for dedicated workgroups or users. Databases can be filled directly with arbitrary trait data not limited to those included in the public instance.
3. Application Programming Interface (API): We also provide an open API that allows third-party programs to make calls to the public instance, or if available also local instances.

Extensive documentation on the code, but also tutorials for users and guides for administrators to host local instances and software developers to use the API are available at the GitHub repository, the public instance and https://fennec.readthedocs.io.

### Mapping community and network data to traits

Minimum requirement for using the Fennec is to provide a community or network table, including taxa as rows and samples (communities)/ taxa (networks) as columns. Cells are considered as abundances but also presence/absence data can be used. Beyond this, users may provide own taxonomy data as an alternative to the default NCBI taxonomy (Federhen, 2012)) and meta-data for the samples. These tables may be uploaded separately as tab-separated text- or combined BIOM-files (Ankenbrand et al., 2017; McDonald et al., 2012) (Figure 1). Files can be managed using the project page of Fennec (Figure 2B). Depending on the user input, taxa are mapped using scientific names or common database identifiers (e.g. NCBI-taxonomy-ID, EOL-ID).

Traits to be analyzed can be explored and selected via the web interface, and added as meta-data to the project (Figure 2A). All data available for the selected traits and taxa of interest are automatically linked into the dataset. If multiple values are available for a single trait and organism combination, they are automatically aggregated, i.e. categorical traits are unified and numerical traits are averaged. To make trait usage as transparent and flexible as possible and to facilitate proper attribution along with aggregated trait values, trait citations for individual values are provided alongside actual traits. Citations can be exported as a separate table and are included in any downloaded BIOM file.

After the mapping, communities are enriched with selected meta-data and can be further processed with standard analytical and statistical software. Download options are individual tables or as a single BIOM file including all information (e.g. for fast integration in R with phyloseq (McMurdie et al., 2013) or QIIME (Caporaso et al., 2010)).

To provide basic analytical plots directly in the workbench, we integrated and modified the open-source project Phinch (Bik et al., 2014). This allows quick interactive exploration of species and trait distributions in each sample, groups, or aggregated by trait types (Figure 2D).

### Data for the public instance

A public instance of Fennec is hosted at https://fennec.molecular.eco and freely available for direct usage. Taxonomy data in this instance consists of a full representation of the NCBI Taxonomy database (Federhen, 2012 accessed 21/03/2018, >1.7 million taxa). Currently only a small fraction (about 73,500) of the taxa in the database have associated traits. We aim to extend this set of traits but also allow users to contribute traits to the general public database. A mapping of EOL-IDs (according to https://opendata.eol.org/dataset/hierarchy_entries, accessed on 04/04/2017) has also been imported, so that full-text information about taxa is available where EOL offers such (Parr, Wilson, Leary, et al., 2014). Currently, and as a starting seed, trait data from TraitBank (Parr, Wilson, Schulz, et al., 2014), EPPO (EPPO, 2017), the World Crops Database (Bijlmakers, 2017), the cavity-nesting bees and wasps database (Budrys et al., 2014, part of the SCALES project (Henle et al., 2014)), IUCN (IUCN, 2017), and ProTraits (Brbić et al., 2016) have been imported which is subject to continuous extension. We aim to maintain the high quality of these publicly available traits. The integration of more traits is a steadily ongoing process. While the bulk of trait data is gathered from databases, in the next release users can also participate in the uploading of trait data so that this process can be actively supported by the community.

### Importing organism and trait data

Traits are imported using a simple table containing, at a minimum the organism and trait value, a citation and optionally ontology URL as columns, with each entry as a new row. Two trait formats are currently supported: categorical and numerical. Categorical traits may also include an ontology URL for their value, supporting the hierarchical classification characteristics. Numerical trait types may be uploaded with an associated unit.

Currently users can upload own traits as project-specific meta-data or in local instances accessible for all users if allowed by the administrator. For the public instance, we plan a moderation system for a future release. All data will undergo a limited manual verification which, for example, ensures that units are correctly standardized but does not verify the correctness of the underlying data. Instead, the user-names of uploaders will be permanently linked to the data to be able to address future changes and updates, alongside corresponding citation information.

For local instances, arbitrary organism and trait data can be imported into the Fennec database by the local administrator using the command line interface. Those trait entries are not subject to a central verification process, but available instantly. This allows creation of instances tailored to specific organism groups and associated research questions, with responsibility for local administrators to ensure quality. Either the global NCBI-taxonomy data can be used or custom taxonomy data provided. Each imported organism receives a unique Fennec-ID which can be linked on-the-fly to other identifiers like NCBI-Taxonomy-IDs or EOL-IDs. The linked EOL-ID is used to provide dynamic content for each organism using the EOL-API (Figure 2C).

## Discussion

The Fennec is a useful tool for automated mapping from taxonomic data to functional meta-data of whole communities. This can be done with user-supplied traits or traits data-mined from trait databases. A growing public instance is available for analyses in pollination studies. It can be accessed via a graphical web interface or by calls via an API. Local instances can be tailored to specific organism groups, workgroups and research questions. The workbench provides basic visualization options for mapped data, as well as export options in various formats for use with downstream software.

### Usage examples

The online documentation at https://fennec.readthedocs.io also includes a case study for pollination ecology to demonstrate how the Fennec can be used in practice. We specifically provide a test data set of 384 pollen samples collected by two closely related megachilid solitary bee species that were assessed using next-generation sequencing based metabarcoding (Sickel et al., 2015), and analyzed with the Fennec with respect to plant growth habit. In addition, the species show quantitative and qualitative differences in their choices for crop plants as a pollen source. Furthermore, the tutorial covers screening for foraging on invasive or vulnerable plant species, which are both easily identified using the Fennec enriched dataset and the visualization options. More generally, the Fennec proves useful wherever more than just taxonomic information is important in multiple species assemblages, for example microbiome ecology, species interaction networks, or plant-herbivore analyses. We aim to frequently expand the tutorials and also encourage users to contribute such.

### Outlook and limitations

The main factor restricting FENNEC’S utility is the currently limited amount of trait data available in a usable format. It became apparent while building the Fennec that a lot of trait data is available online, but the majority does not adhere to the FAIR principles (Findable, Accessible, Interoperable, Reusable) (Mons et al., 2017; Wilkinson et al., 2016). Also, licensing of the data is a common problem; data can only be efficiently re-used if it is open and citable in addition to being FAIR (Katz, 2017). It is essential to guarantee trait data collectors that data re-users are able to properly cite all data sources, e.g. by adhering to the FORCE11 data citation principles (Martone, 2014). Fennec supports this by preserving all relevant information. We therefore encourage trait data collectors to make their data available via existing platforms like TraitBank (Parr, Wilson, Schulz, et al., 2014), and thereby also usable for downstream analysis tools like Fennec and ultimately available to the whole research community.

## Conclusion

Fennec as a tool provides valuable assistance to analyze ecological data with more than just taxonomy. Species traits and meta-data like threat status and economic importance help to answer questions beyond the mere presence or absence of organisms. The public instance can be used as a reference, to try features of Fennec and analyze datasets with data from other public databases. Self-tailored local instances with private data increases the range of applications.

The main current limitation is trait data availability, but we expect an increase with time, ongoing open-science initiatives and scientific journals requiring public data deposition before publication. Besides developing a public automatic mapping procedure, we also aim to demonstrate the importance of making trait data publicly available and its usefulness in follow up studies. We thus advocate the submission of trait data to public databases.

Despite limitations in data availability, the Fennec is technically already able to enrich ecological community analyses with trait information. Its usefulness is expected to increase due to continued development guided by user feedback, integration of more analysis tools, better taxonomic resolution, and increasing availability of suitable trait data.

## Acknowledgements

This study was supported by the German Excellence Initiative to the Graduate School of Life Sciences, University of Würzburg (405/1517) to MJA and the DFG (KE1743/7-1) to AK. We are grateful to the EOL for quick, open and helpful responses to our emails. Many thanks to A. Voulgari-Kokota, N. Terhoeven, J. Schultz and A. Claßen for beta testing, suggestions and valuable feedback.

## Data Accessibility

The source code of Fennec including documentation is available at https://github.com/molbiodiv/fennec. This source code is also archived at Zenodo with doi: 10.5281/zenodo.591305.

## Author Contributions

MJA and AK conceived the project, as well as designed methodology and software; MJA and SH wrote the code with support by FF and LW; MJA analyzed the case study and drafted the manuscript. All authors contributed critically to the drafts and gave final approval for publication.

